# Multi-locus analysis of genomic time series data from experimental evolution

**DOI:** 10.1101/006734

**Authors:** Jonathan Terhorst, Yun S. Song

## Abstract

Genomic time series data generated by evolve-and-resequence (E&R) experiments offer a powerful window into the mechanisms that drive evolution. However, standard population genetic inference procedures do not account for sampling serially over time, and new methods are needed to make full use of modern experimental evolution data. To address this problem, we develop a Gaussian process approximation to the multi-locus Wright-Fisher process with selection over a time course of tens of generations. The mean and covariance structure of the Gaussian process are obtained by computing the corresponding moments in discrete-time Wright-Fisher models conditioned on the presence of a linked selected site. This enables our method to account for the effects of linkage and selection, both along the genome and across sampled time points, in an approximate but principled manner. Using simulated data, we demonstrate the power of our method to correctly detect, locate and estimate the fitness of a selected allele from among several linked sites. We also study how this power changes for different values of selection strength, initial haplotypic diversity, population size, sampling frequency, experimental duration, number of replicates, and sequencing coverage depth. In addition to providing quantitative estimates of selection parameters from experimental evolution data, our model can be used by practitioners to design E&R experiments with requisite power. Finally, we explore how our likelihood-based approach can be used to infer other model parameters, including effective population size and recombination rate, and discuss extensions to more complex models.

## Introduction

A common study design in population genetics consists of collecting genomic variation data from living organisms to make inferences about unobserved evolutionary and biological phenomena. The many areas where this design has been applied include demographic inference (see [1] for a recent review), recombination rate estimation [2–6], and detection of natural selection [7–13]. Recently, there has been much interest in utilizing time series genetic data—e.g., from ancient DNA [14–19], experimental evolution of a population under controlled laboratory environments [20–24], or direct measurements in fast evolving populations [25]—to enhance our ability to probe into evolution. In particular, understanding the genetic basis of adaptation to changes in the environment can be significantly facilitated by such temporal data. Specifically, the dynamics of allele frequencies in an evolving population potentially convey added information about how the genome functions [26], information which is inaccessible to methods which operate only on a static snapshot of that genome.

An experimental methodology which serially interrogates the genomes of an controlled population over time could potentially yield new insights. In fact, this methodology can now be realized thanks to the advent of next-generation sequencing. By sequencing successive generations of model organisms raised in a controlled environment, genetic time series data can be generated which describe evolution at nucleotide resolution [23, 26–28]. This so-called evolve-and-resequence (henceforth, E&R) methodology is fundamentally different than the observational approach described above, and new inference procedures are needed to analyze this type of data.

In this work we present such a procedure and study its ability to perform a number of testing and estimation tasks relevant to population genetics. Our method is based on an approximation to the multi-locus Wright-Fisher process, and is well-suited to the small population, discrete generation, and random mating setting in which many E&R experiments are conducted. Furthermore, because it is based on a canonical population genetic model of genome evolution, our method can directly estimate population genetic quantities such as fitness, dominance, recombination rate, and effective population size. It can also be used to design future experiments with sufficient power to reliably infer these quantities.

### Related work

There is a small but growing literature on the analysis of evolve-and-resequence data. Feder *et al.* [29] present a statistical test for detecting selection at a single biallelic locus in time series data. (Although it is not a major focus, their method can also be used to estimate the selection parameter.) Similar to our method, they model the sample paths of the Wright-Fisher process as Gaussian perturbations around a deterministic trajectory in order to obtain a computable test statistic. However, their aim is slightly different from ours in that they analyze yeast and bacteria data sets where the population size is both large and must be estimated from data. Here we focus on population sizes which are smaller and more typical of experiments performed on higher organisms, for example mice or *Drosophila*. We generally assume that the effective population size is known but also test our ability to estimate it from data. Also, because of the increased amount of drift present in the small population regime, we necessarily restrict our attention to selection coefficients which are somewhat larger than those considered by Feder *et al.* Finally, although Feder *et al.* do study the performance of their method when time series data are corrupted by noise due to finite sampling (as in e.g. a next-generation sequencing experiment), they do not model this effect. Here we properly account for the effect of sampling by integrating over the latent space of population-level frequencies when computing the likelihood.

Another related work is Baldwin-Brown *et al.* [30], which presents a thorough study of the effects of sequencing effort, replicate count, strength of selection, and other parameters on the power to detect and localize a single selected locus segregating in a 1 Mb region. Results are obtained by simulating data under different experimental conditions and comparing the resulting distributions of allele trajectories under selection and neutrality using a modified form of *t*-test. Because it is not model-based, this method is incapable of performing parameter estimation. As a result of their study, Baldwin-Brown *et al.* present a number of design recommendations to experimenters seeking to attain a given level of power to detect selection. In a related work, Kofler and Schlötterer [31] carried out forward simulations of whole genomes to provide guidelines for designing E&R experiments to maximize the power to detect selected variants.

Illingworth *et al.* [32] derive a probabilistic model for time series data generated from large, asexually reproducing populations. The population size is sufficiently large (on the order of *∼* 10^8^) that population allele frequencies evolve quasi-deterministically. The deterministic trajectories are governed by a system of differential equations describing the effect of a selected (“driver”) mutation on nearby linked neutral (“passenger”) mutations. Randomness arises due to the finite sampling of alleles by sequencing. The main difference between the setting of Illingworth *et al.*’s and our own concerns genetic drift. While drift may be ignored when studying a large population of microorganisms, we show that it confounds our ability to detect and estimate selection in populations of order *∼* 10^3^. Thus, for E&R studies on (smaller) populations of macroscopic organisms, methods which assume that allele frequencies evolve deterministically may not perform as well as those which explicitly take drift into account.

Topa *et al.* [33] present a Bayesian model for single-locus time series data obtained by next-generation sequencing. In each time period, the allele count is modeled as a draw from a binomial distribution with number of trials equal to the depth of sequencer coverage, and success probability equaling the population-level allele frequency. The posterior allele frequency distribution is used to test for selection by comparing a neutral model to one in which unobserved allele frequencies to depend on time. In the non-neutral case, a Gaussian process is used to allow for directional selection acting on the posterior allele frequency distributions.

Finally, Lynch *et al.* [34] derive a likelihood-based method for estimating population allele frequency at a single locus in pooled sequencing data. The method allows for the possibility of sequencing errors as well as subsampling the population prior to sequencing. Using theoretical results as well as simulations, the authors give guidelines on the (subsampled) population size and coverage depth needed to reliably detect a difference in allele frequency between two populations. Unlike the other methods surveyed here, the approach of Lynch *et al.* is not designed to analyze time series data. Hence the data requirements needed to reliably detect allele frequency changes using their method—for example, sequencing coverage depth of at least 100 reads—are potentially greater than for methods are informed by a population-genetic model of genome evolution over time.

### Novelty of our method

Our method differs from the above-mentioned approaches in several regards. To the best of our knowledge, ours is the first method capable of analyzing time series data from multiple linked sites jointly. As we show below, this is advantageous when studying selection in E&R data. Furthermore, it enables us to analyze features of these data which cannot be studied using single-locus models, such as local levels of linkage disequilibrium and the effect of a recombination hotspot. Additionally, because our model is based on a principled approximation to the Wright-Fisher process, it can numerically estimate the selection coefficient, dominance parameter, recombination rates, and other population genetic quantities of interest. In this way it is distinct from the aforementioned simulation-based methods [30, 31], methods which only focus on testing for selection [29, 30, 33], or methods based on general statistical procedures which are not specific to population genetics [33,34].

### Software availability

An open-source software package implementing the method described in this paper will be made publicly available.

## Results

We tested our method on simulated data designed to capture the essential features of an E&R experiment. See **Methods** for the details on simulation. Briefly, it consisted of cloning a set of *F* homozygous founder lines (whose haplotypes are assumed to be known) to form an experimental population of *N* diploid organisms, which were then simulated forwards in time for *T* generations according to the Wright-Fisher random mating model. For each segregating site, we assumed that there are two alleles, denoted *A*_0_ and *A*_1_. The experiment was repeated using the same starting conditions to form *R* experimental replicates. After the simulation terminated, the frequency of allele *A*_1_ was recorded for each combination of segregating site, time period and replicate, possibly with introduced sampling error; this setup mimics pooled sequencing. The input to the model consisted of these time series allele frequency data along with the haplotypes of the founder lines.

Certain aspects of the simulation were varied to test different aspects of the model; these changes are described more fully in their respective sections below. Unless otherwise noted, the simulations were performed using *F* = 10 founder lines, census population size *N* = 1000, sampling at generations *t_i_* ∈ {10, 20, 30, 40, 50}, *R* = 3 experimental replicates and a region of size *L* = 10^5^ sites. These values were chosen to reflect a typical E&R experiment and we refer to them in the sequel as the “default” parameter values. Sequencing coverage depth is denoted by *C*, with *C* = *∞* corresponding to having perfect knowledge of the population allele frequencies. We use *C* = *∞* in the default parameter setting to upper bound the performance of our method, but also consider *C* = 10 and 30 to investigate the effect of uncertainty in allele frequency estimation.

A common objective in E&R experiments is to detect genetic adaptation. For example, a population may be partitioned, with one subgroup placed in a new environment. Upon running an E&R experiment, one wishes to 1) determine whether a fitness difference exists between the control and subject groups; 2) find the alleles responsible for the adaptation; and 3) estimate the strength of selection acting on these alleles. To test our model’s ability to perform each of these tasks, we simulated E&R experiments in which a segregating site in the founding population was chosen uniformly at random and placed under selection. The relative fitnesses of *A*_0_*/A*_0_ and *A*_1_*/A*_1_ homozygote genotypes are respectively given by 1 and 1 + *s*, while the relative fitness of the heterozygote *A*_0_*/A*_1_ is 1 + *hs*. In what follows, we assume *h* = 1*/*2 unless stated otherwise.

### Testing for selection

Let *s_i_* denote the coefficient of selection at segregating site *i* = 1, …, *K*, where *K* is the total number of segregating sites in the region being considered. We wish to test the following null and alternative hypotheses:

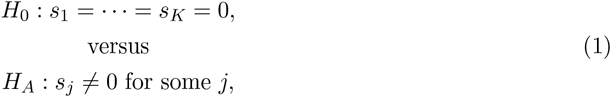

which can be implemented using a standard likelihood ratio test. As the number *R* of experimental replicates grows large, the distribution of the test statistic under the null hypothesis tends to a *χ*^2^ distribution. However, since *R* was set to a realistic (i.e., small) value in our experiments, we found that the test performed better if the null distribution was determined empirically by performing additional simulations under neutrality.

Using the default parameter settings mentioned earlier, Figure 1 displays the test’s receiver operating characteristic (ROC) curve for various strengths *s* of selection. Each curve was estimated from 1000 simulations. Strong selection (*s* = 0.1) is easily distinguished from neutrality, with all cases detected at a false positive rate of only 0.5%. Moderate selection (*s* = 0.05) is more challenging to detect, but the test still has 90% power to detect selection with a false positive rate of 5.8%. Weaker selection (*s* = 0.02) is more challenging still; achieving 50% power in this case would entail a false positive rate of 28%.

**Figure 1.**
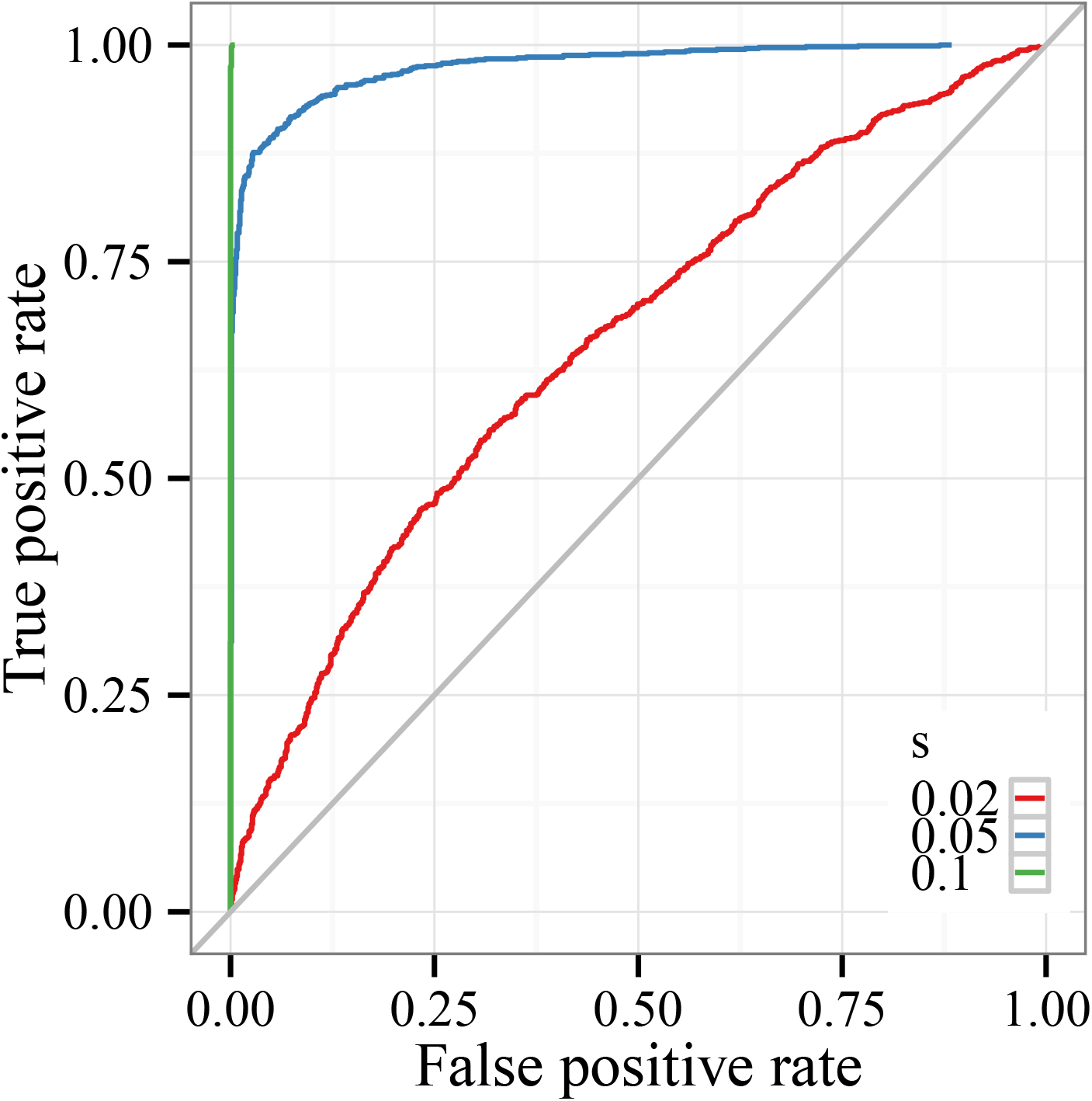
Receiver operating characteristics (ROC) when testing for selection in a region under the default parameter setting. Each ROC curve was estimated using 1000 simulations. For each selection regime, the curve was calculated by comparing the distribution of the maximum likelihood ratio over all segregating sites in a region of length 100 kb with the distribution of the same statistic under neutrality. As the figure shows, strong selection (*s* = 0.1) is easy to distinguish from neutrality, with a negligible false positive rate. Moderate selection (*s* = 0.05) is more challenging to detect, but the test still has 90% power with a false positive rate of *∼* 6%. Weaker selection (*s* = 0.02) poses more challenge; in this case achieving 50% power would entail a false positive rate of 28%.

Weaker selection is harder to detect because it is difficult to distinguish from drift. Thus, one option for improving sensitivity to weaker selection is to increase the population size used in the experiment. To study how drift influences our ability to detect weaker selection, we ran additional simulations with larger population sizes *N* ∈ {2000, 5000} while holding the remaining experimental parameters fixed. Results from these experiments are shown in Figure 2. Selection at the *s* = 0.05, 0.10 levels can be detected almost without error for these larger population sizes. Though weaker selection remains difficult to detect, our method is able to detect selection at strength *s* = 0.02 with 75% precision (*N* = 2000) and 90% precision (*N* = 5000) while maintaining a false positive rate of 8%.

**Figure 2.**
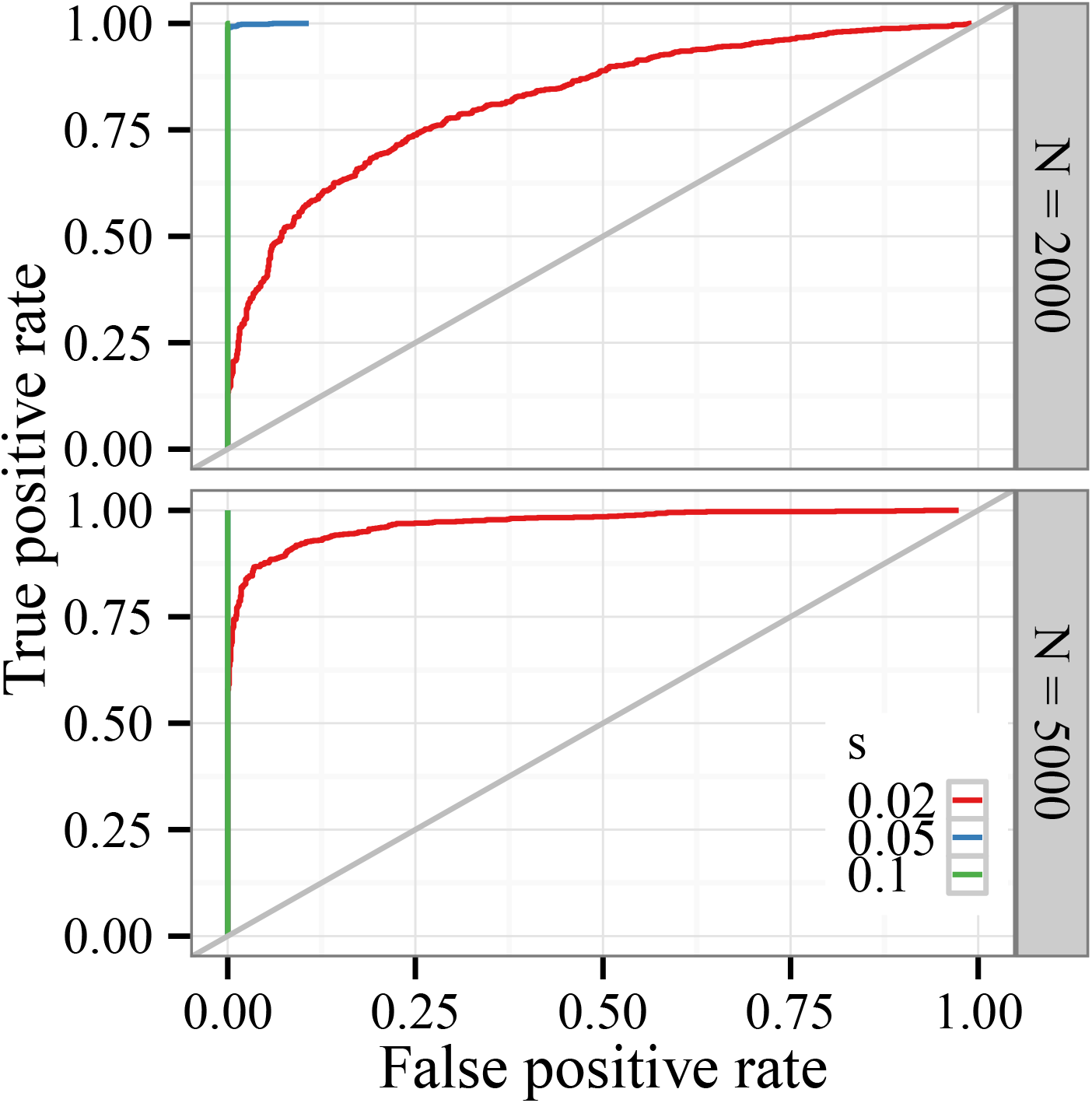
Receiver operating characteristics when testing for selection in E&R experiments with larger population sizes. Parameters for each simulation were the same as in Figure 1, except that the population size was increased to *N* = 2000 (top panel) and *N* = 5000 (bottom panel). Comparing these ROC curves with those in Figure 1, we see that increasing the population size by only a few folds significantly improves the performance of the test for selection.

### Locating the selected site

Once selection has been detected in a region, it is desirable to map the selected site as accurately as possible. An obvious estimator in this case is to declare the site with the highest log-likelihood ratio (versus a neutral model) from the preceding test to be the selected site. Table 1 shows how this estimation procedure performed for different strengths of selection. Since increased haplotypic diversity should make it easier to localize the site responsible for non-neutral behavior, we performed further simulations where the number of founder haplotypes was increased to *F* = 20 and 30. Two measures of the accuracy are displayed. The first set of columns examines the distribution of the distance (in base pairs) between the estimated and true selected site. As expected, selection becomes easier to localize as it becomes stronger and as the number of founder haplotypes grows. With strong selection and 20 or more founder haplotypes (bottom two rows), the method correctly pinpointed the exact location of the selected site in over 50% of the simulations. The top rows of Table 1 indicate that weaker selection (*s* = 0.02) is difficult to localize precisely using this method; the median estimated distance from the true selected site was 25-30 kb in these cases.

**Table 1.**
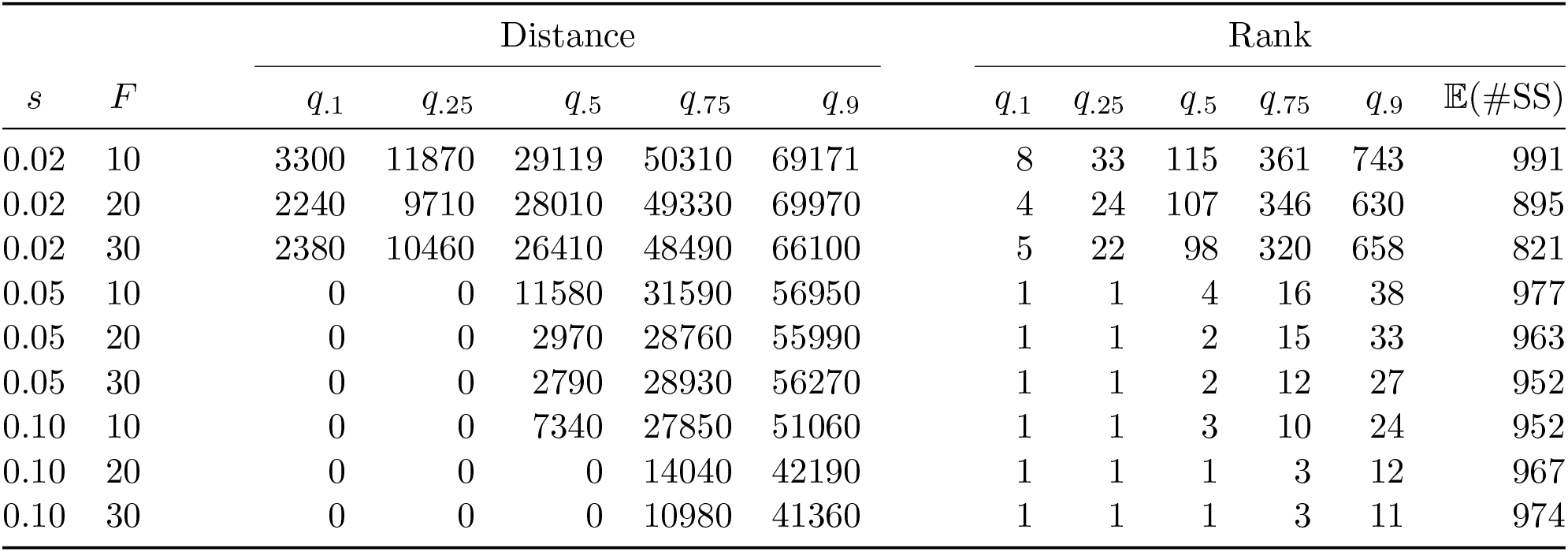
Results of localization procedure. The two sets of columns display percentiles of the distance in base pairs from the estimated selected site to the true selected site, and of the average rank (in terms of likelihood ratio) of the true selected site. The column labeled *q_j_* corresponds to the *j*th percentile. The column labeled 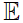(#SS) shows the average number of segregating sites observed over all simulations.

| $s$ | $F$ | Distance | | | | | Rank | | | | | $\mathbb{E}(\#SS)$ |
| --- | --- | --- | --- | --- | --- | --- | --- | --- | --- | --- | --- | --- |
| | | $q.1$ | $q.25$ | $q.5$ | $q.75$ | $q.9$ | $q.1$ | $q.25$ | $q.5$ | $q.75$ | $q.9$ | |
| 0.02 | 10 | 3300 | 11870 | 29119 | 50310 | 69171 | 8 | 33 | 115 | 361 | 743 | 991 |
| 0.02 | 20 | 2240 | 9710 | 28010 | 49330 | 69970 | 4 | 24 | 107 | 346 | 630 | 895 |
| 0.02 | 30 | 2380 | 10460 | 26410 | 48490 | 66100 | 5 | 22 | 98 | 320 | 658 | 821 |
| 0.05 | 10 | 0 | 0 | 11580 | 31590 | 56950 | 1 | 1 | 4 | 16 | 38 | 977 |
| 0.05 | 20 | 0 | 0 | 2970 | 28760 | 55990 | 1 | 1 | 2 | 15 | 33 | 963 |
| 0.05 | 30 | 0 | 0 | 2790 | 28930 | 56270 | 1 | 1 | 2 | 12 | 27 | 952 |
| 0.10 | 10 | 0 | 0 | 7340 | 27850 | 51060 | 1 | 1 | 3 | 10 | 24 | 952 |
| 0.10 | 20 | 0 | 0 | 0 | 14040 | 42190 | 1 | 1 | 1 | 3 | 12 | 967 |
| 0.10 | 30 | 0 | 0 | 0 | 10980 | 41360 | 1 | 1 | 1 | 3 | 11 | 974 |

When running these experiments we observed that the likelihood ratio of the true selected site was often just beneath that of the site with the maximum likelihood ratio. Rather than estimating the single site most likely to be driving selection, the method could also be used to provide a list of candidate sites which could then be investigated using additional knowledge about, for example, the functional effect of each mutation. The second set of columns in Table 1 examines the distribution of the rank of the true selected site when all segregating sites in the region are sorted according to their likelihood ratio. For medium and strong selection the true selected site was among the top five out of at least 800 segregating sites in at least half of the simulations, and within the top 40 in over 90% of them. With weaker selection the situation is again more difficult; in over half the simulations there were at least 100 other sites which were as likely to be under selection as the true site.

We also studied how coverage depth affects the ability to map the selected site. For *F* = 10, Table 2 repeats the analysis of Table 1 when the data are sampled at simulated coverage depths of 10 and 30 short-reads. Comparing the two tables, we see that the additional noise introduced by sequencing makes the problem of localizing the selected site more difficult; the modal estimate is often separated from the true site by tens of kilobases. Nevertheless, in more than half the trials performed we observed that a strongly selected site would be among the top ten segregating sites (in terms of likelihood ratio; see Table 2, last two rows). For medium selection, increasing coverage depth from 10 to 30 short-reads improved our ability to map the selected site by several kilobases, and more than halved the number of segregating sites we would need to examine before encountering the selected site. Weaker selection, already difficult to detect without sampling, is even more so when noise is introduced.

**Table 2.** Results of localization procedure for the case of *F* = 10 and finite coverage depth. Data were generated as in Table 1 and then sampled to simulate sequencing. Coverage depth is indicated in the column labeled *C*. The column labeled *q_j_* corresponds to the *j*th percentile. The column labeled 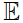(#SS) shows the average number of segregating sites observed over all simulations.

| $s$ | $C$ | Distance | | | | | Rank | | | | | $\mathbb{E}(\#SS)$ |
| --- | --- | --- | --- | --- | --- | --- | --- | --- | --- | --- | --- | --- |
| | | $q.1$ | $q.25$ | $q.5$ | $q.75$ | $q.9$ | $q.1$ | $q.25$ | $q.5$ | $q.75$ | $q.9$ | |
| 0.02 | 10 | 3480 | 11431 | 28360 | 49930 | 70050 | 29 | 99 | 283 | 615 | 853 | 988 |
| 0.02 | 30 | 4040 | 12260 | 28370 | 50010 | 68360 | 24 | 78 | 225 | 514 | 871 | 967 |
| 0.05 | 10 | 590 | 8100 | 23590 | 46120 | 67040 | 2 | 8 | 35 | 109 | 262 | 958 |
| 0.05 | 30 | 60 | 5900 | 20810 | 41260 | 63341 | 2 | 5 | 18 | 45 | 110 | 950 |
| 0.10 | 10 | 0 | 1210 | 16670 | 39840 | 60680 | 1 | 2 | 8 | 23 | 44 | 872 |
| 0.10 | 30 | 0 | 150 | 14530 | 36701 | 58210 | 1 | 2 | 4 | 17 | 32 | 837 |

### Estimating the strength of selection

Once a selected site has been located, it is desirable to numerically quantify the fitness of the *A*_1_ allele. Table 3 describes the distribution of these estimates for various combinations of selective strength, coverage depth, and model complexity (i.e., the number of loci in the Gaussian process approximation). For each of the simulations above we estimated *s* by maximum likelihood. To separate the ability of our model to estimate selection from its ability to locate the selected site, we assumed that the selected site was already known when performing these estimates. Aside from varying selection strength, we also examined how coverage depth and the number of loci used for estimation affected the quality of the estimates. For each parameter combination, the table displays the mean, median and inter-quartile range (IQR) of the distribution of the maximum likelihood estimate ŝ of s.

**Table 3.**
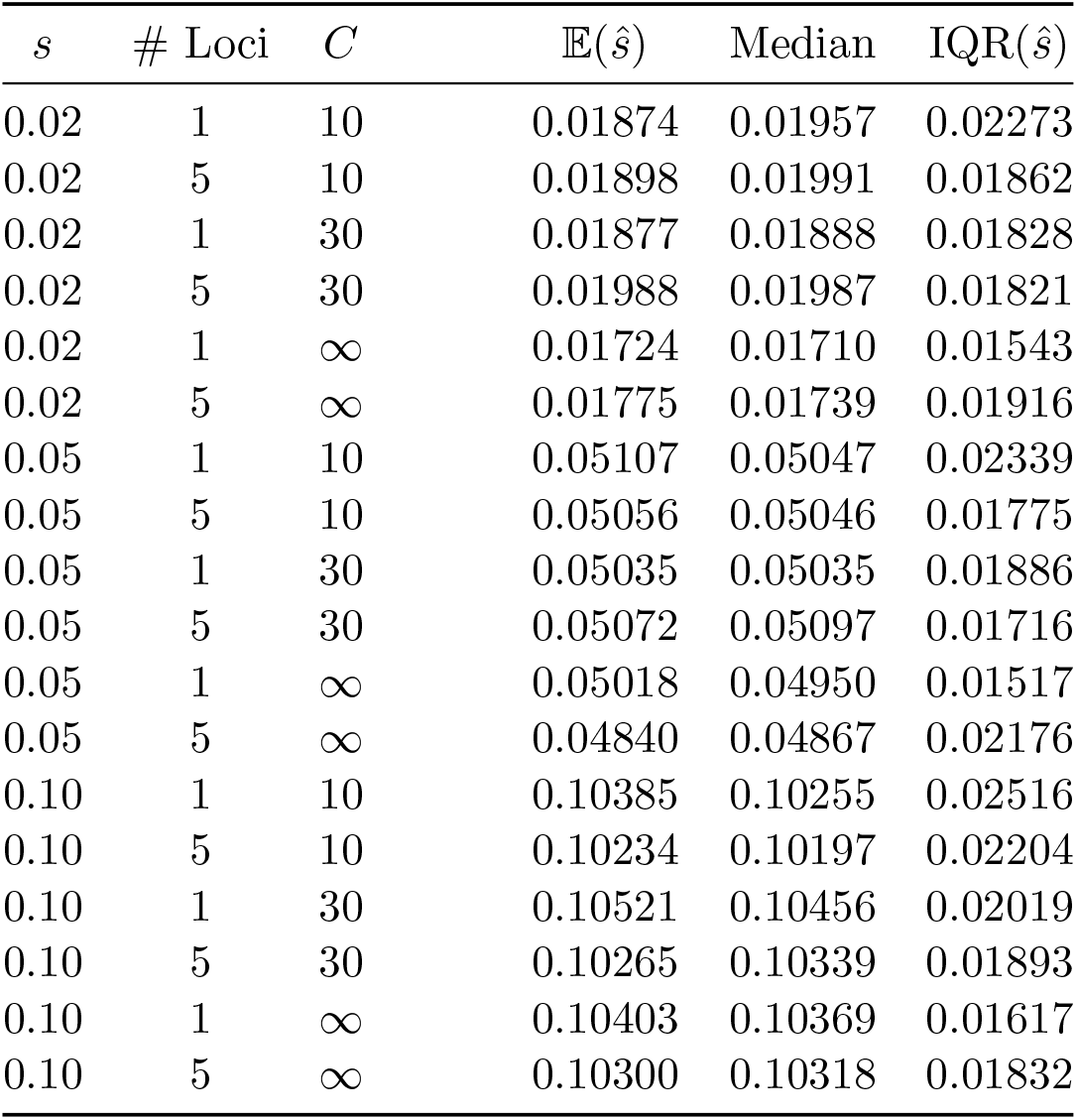
Estimation of selection coefficient. For each combination of selection strength, model complexity, and coverage depth (*s*, # Loci, and *C*, respectively), the rightmost columns display the average, median and inter-quartile range of the selection estimate ŝ obtained from 1000 simulations. Rows with *C* = *∞* denote simulations when the population-level allele frequency was known without error.

| $s$ | # Loci | $C$ | $\mathbb{E}(\hat{s})$ | Median | IQR($\hat{s}$) |
| --- | --- | --- | --- | --- | --- |
| 0.02 | 1 | 10 | 0.01874 | 0.01957 | 0.02273 |
| 0.02 | 5 | 10 | 0.01898 | 0.01991 | 0.01862 |
| 0.02 | 1 | 30 | 0.01877 | 0.01888 | 0.01828 |
| 0.02 | 5 | 30 | 0.01988 | 0.01987 | 0.01821 |
| 0.02 | 1 | ∞ | 0.01724 | 0.01710 | 0.01543 |
| 0.02 | 5 | ∞ | 0.01775 | 0.01739 | 0.01916 |
| 0.05 | 1 | 10 | 0.05107 | 0.05047 | 0.02339 |
| 0.05 | 5 | 10 | 0.05056 | 0.05046 | 0.01775 |
| 0.05 | 1 | 30 | 0.05035 | 0.05035 | 0.01886 |
| 0.05 | 5 | 30 | 0.05072 | 0.05097 | 0.01716 |
| 0.05 | 1 | ∞ | 0.05018 | 0.04950 | 0.01517 |
| 0.05 | 5 | ∞ | 0.04840 | 0.04867 | 0.02176 |
| 0.10 | 1 | 10 | 0.10385 | 0.10255 | 0.02516 |
| 0.10 | 5 | 10 | 0.10234 | 0.10197 | 0.02204 |
| 0.10 | 1 | 30 | 0.10521 | 0.10456 | 0.02019 |
| 0.10 | 5 | 30 | 0.10265 | 0.10339 | 0.01893 |
| 0.10 | 1 | ∞ | 0.10403 | 0.10369 | 0.01617 |
| 0.10 | 5 | ∞ | 0.10300 | 0.10318 | 0.01832 |

Several interesting features emerge from the table. Inter-quartile range is of roughly the same order across scenarios, so that estimation error shrinks relatively as selection become stronger. For one-locus models, IQR shrinks as coverage depth increases. For multi-locus models the effect of increasing the number of sites used to perform estimation is interesting. When the data are observed without noise, we saw little improvement in the accuracy of *ŝ* when using a single-locus model fit only to data from the selected site versus a multi-locus model which also took the trajectories of linked sites into account. In fact, in several cases this cause the estimates to become more dispersed as the trajectory of the selected allele had relatively less weight in the likelihood calculation. On the other hand, when allele frequencies are sampled with noise we see that estimates of *ŝ* obtained from a five-locus model generally have smaller IQR, particularly in the low-coverage-depth case *C* = 10. These findings are confirmed in Figure 3, which displays density estimates for the residual *s − ŝ* for each of these cases presented in the table. Compared with the one-locus model, the five-locus model which takes additional data from linked sites into account produces estimates which are more concentrated around the true parameter value. Thus, when the data are noisy (i.e., when *C* is small), the trajectories of nearby linked sites provide useful information concerning the (unobserved) population frequency of the selected allele as it evolves over time.

**Figure 3.**
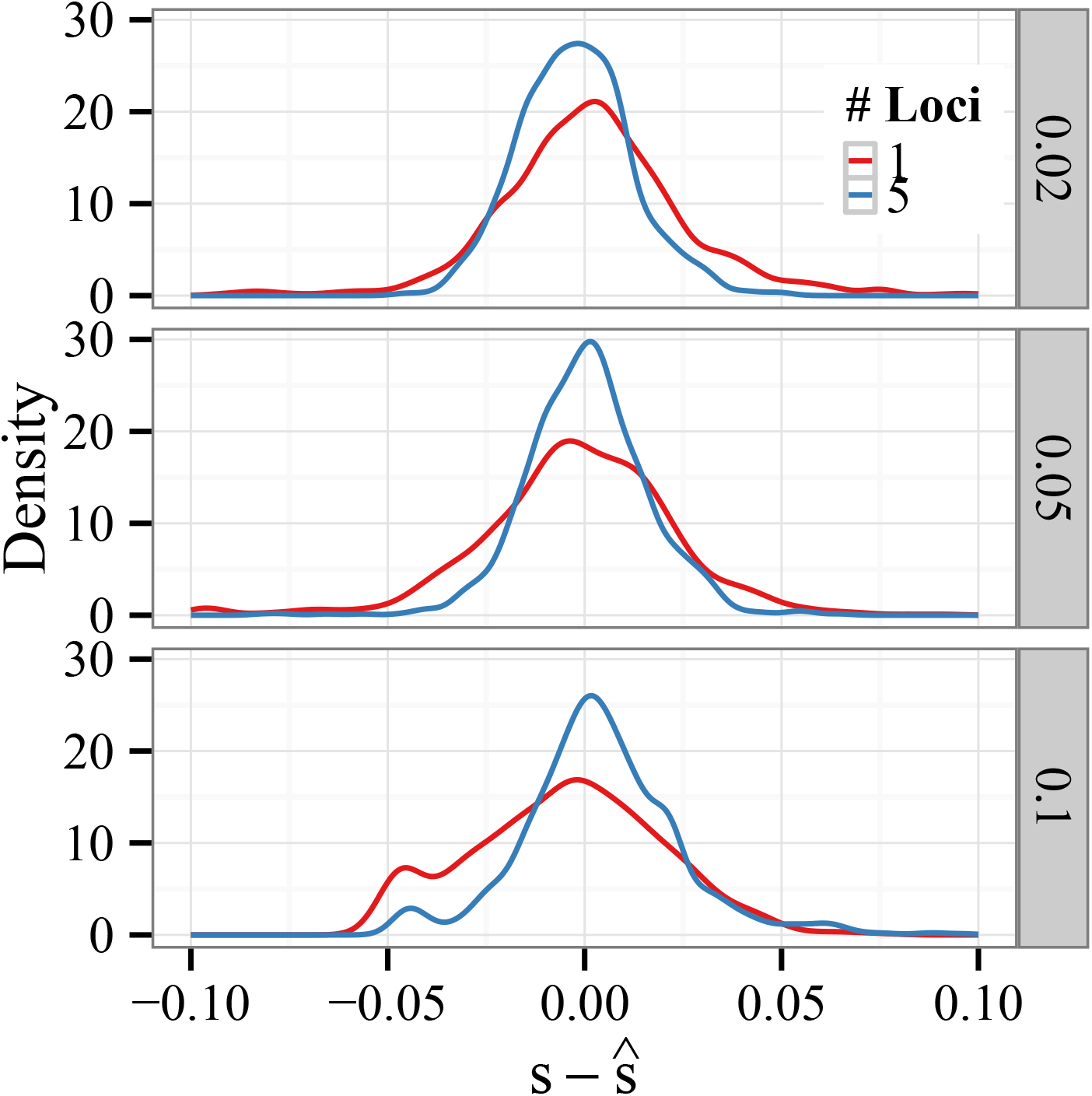
Estimated error density of with sampling. Data were generated using the standard parameters and sampled to a depth of 10 reads per site. Density estimates for the residual *s* − *ŝ* for *s* = 0.02, 0.05, 0.10 (top to bottom) are plotted. The red and green lines denote the density estimates obtained using one-and five-locus models, respectively. The five-locus model, which takes additional data from linked sites into account, produces estimates which are more concentrated around the true parameter value.

We observed a slight negative bias for weaker selection and a slight positive bias for medium and strong selection, which can be attributed to loss or fixation of the selected allele. Indeed, estimated selection may be negative when a weakly selected allele segregating at low frequency is lost due to drift; similarly, there is a tendency to overestimate the strength of selection acting on a high-frequency allele which fixes quickly.

It is also interesting to consider the effect of study design on estimation accuracy. In Table 4 we examine how parameter estimates are affected by sequencing effort and experimental duration. We focus on the limited-coverage case (*C* = 10) since it is most sensitive to adding or removing sequence data from additional generations. For ease of comparison, the first set of rows reproduces data from Table 4, where generations {10, 20, 30, 40, 50} were sequenced. The next subsection examines the case when sequencing effort is reduced to two time periods {25, 50}. The final subsection studies estimation quality when the experimental duration is halved, and only one round of sequencing is performed at generation 25. In all cases we see that the estimators are approximately unbiased, 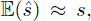 but that their dispersion about the true parameter value is greatly affected by data availability. Sampling genomic data at just a single time period *t* = 25 roughly doubles the IQR of the estimator in each case. Interestingly, with two time periods (*t* ∈ {25, 50}) performance is improved, and the estimator is only somewhat less precise than when sampling at every tenth generation. Finally, as in the previous table we see again that, at least for data sampled at low coverage, estimation performance is unilaterally improved by fitting a multi-locus model versus a single-locus model.

**Table 4.** Effect of sampling frequency on selection coefficient estimation. Column definitions are the same as in Table 3. The three sections correspond to sampling at generations (10, 20, 30, 40, 50), 25, and (25, 50) respectively.

| $s$ | # Loci | $C$ | $\mathbb{E}(\hat{s})$ | Median | $\text{IQR}(\hat{s})$ |
| --- | --- | --- | --- | --- | --- |
| $t_i \in \{10, 20, 30, 40, 50\}$ | | | | | |
| 0.02 | 1 | 10 | 0.01874 | 0.01957 | 0.02273 |
| 0.02 | 5 | 10 | 0.01898 | 0.01991 | 0.01862 |
| 0.05 | 1 | 10 | 0.05107 | 0.05047 | 0.02339 |
| 0.05 | 5 | 10 | 0.05056 | 0.05046 | 0.01775 |
| 0.10 | 1 | 10 | 0.10385 | 0.10255 | 0.02516 |
| 0.10 | 5 | 10 | 0.10234 | 0.10197 | 0.02204 |
| $t_i \in \{25\}$ | | | | | |
| 0.02 | 1 | 10 | 0.01742 | 0.02231 | 0.05067 |
| 0.02 | 5 | 10 | 0.01938 | 0.02086 | 0.03450 |
| 0.05 | 1 | 10 | 0.04958 | 0.04813 | 0.05762 |
| 0.05 | 5 | 10 | 0.04864 | 0.04887 | 0.03045 |
| 0.10 | 1 | 10 | 0.09913 | 0.10167 | 0.05164 |
| 0.10 | 5 | 10 | 0.09930 | 0.09948 | 0.03535 |
| $t_i \in \{25, 50\}$ | | | | | |
| 0.02 | 1 | 10 | 0.01912 | 0.01886 | 0.02799 |
| 0.02 | 5 | 10 | 0.01948 | 0.01953 | 0.01923 |
| 0.05 | 1 | 10 | 0.05149 | 0.05047 | 0.02591 |
| 0.05 | 5 | 10 | 0.05142 | 0.05037 | 0.01969 |
| 0.10 | 1 | 10 | 0.10360 | 0.10256 | 0.03049 |
| 0.10 | 5 | 10 | 0.10139 | 0.10105 | 0.02208 |

### Overdominance estimation

In the preceding discussion, the dominance parameter was fixed at *h* = 1*/*2, so that selection acted additively. Our method is capable of handling general diploid selection. In our experiment, we tested our method’s ability to estimate the effect of overdominance, in which case heterozygotes are fitter than either homozygote. We simulated populations under the conditions *h >*1 and *s* ≪ 1 such that heterozygotes had a relative fitness of 1 + *hs* where *hs* ∈ {0.02, 0.05, 0.10}. Thus, heterozygotes have a fitness advantage of the same order as that which we were able to detect in the additive case.

Results for jointly estimating *h* and *s* are shown in Table 5. A fixed value of *s* = 0.01 was used for fitness in all cases, while *h* was varied. We found that estimating overdominance is difficult when both alleles are initially segregating near their limiting frequency of 1/2, since the resulting allele trajectories appear very similar to those generated by a neutral model with drift. The results in the table are therefore conditioned on the initial allele frequency residing outside of the interval [0.4, 0.6].

**Table 5.**
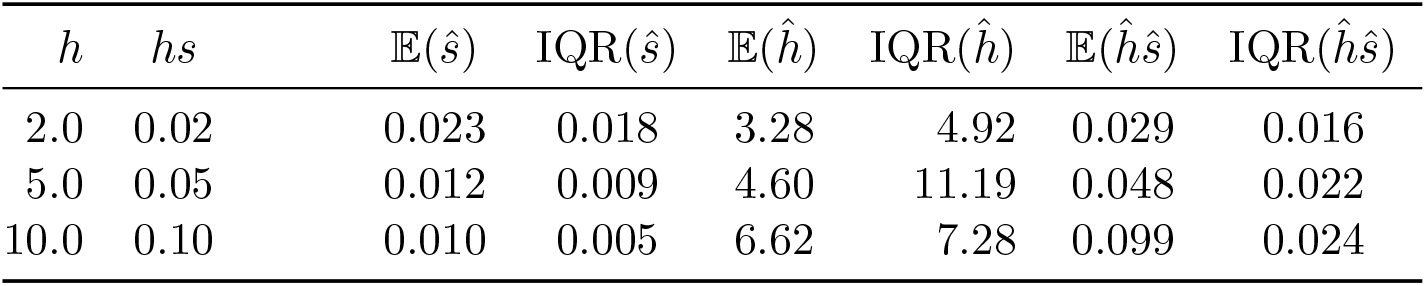
Overdominance estimation. The selection coefficient was fixed at *s* = 0.01 while the dominance parameter *h* was varied. In each simulation, the initial allelic frequency was restricted to lie outside the interval [0.4, 0.6] (see discussion in text).

| $h$ | $hs$ | $\mathbb{E}(\hat{s})$ | $\text{IQR}(\hat{s})$ | $\mathbb{E}(\hat{h})$ | $\text{IQR}(\hat{h})$ | $\mathbb{E}(\hat{h}\hat{s})$ | $\text{IQR}(\hat{h}\hat{s})$ |
| --- | --- | --- | --- | --- | --- | --- | --- |
| 2.0 | 0.02 | 0.023 | 0.018 | 3.28 | 4.92 | 0.029 | 0.016 |
| 5.0 | 0.05 | 0.012 | 0.009 | 4.60 | 11.19 | 0.048 | 0.022 |
| 10.0 | 0.10 | 0.010 | 0.005 | 6.62 | 7.28 | 0.099 | 0.024 |

When considered individually, the estimators *ĥ* and *ŝ* are highly variable (see Table 5, columns 3–6). This behavior is expected since, as witnessed in the previous sections, small values in *s* (specifically, *s* = 0.01) are difficult to detect in experimental data. Encouragingly, a different picture emerges when we consider the product estimator *ĥ* · *ŝ* (see Table 5, columns 7–8). The estimator is close in expectation to the true value *hs* (column 2) and also more tightly concentrated around that value. Density estimates of the product estimator *ĥŝ* are shown in Figure 4 and confirm this finding. Each density estimate has a mode at the true parameter value *hs* and is reasonably concentrated around that value.

**Figure 4.**
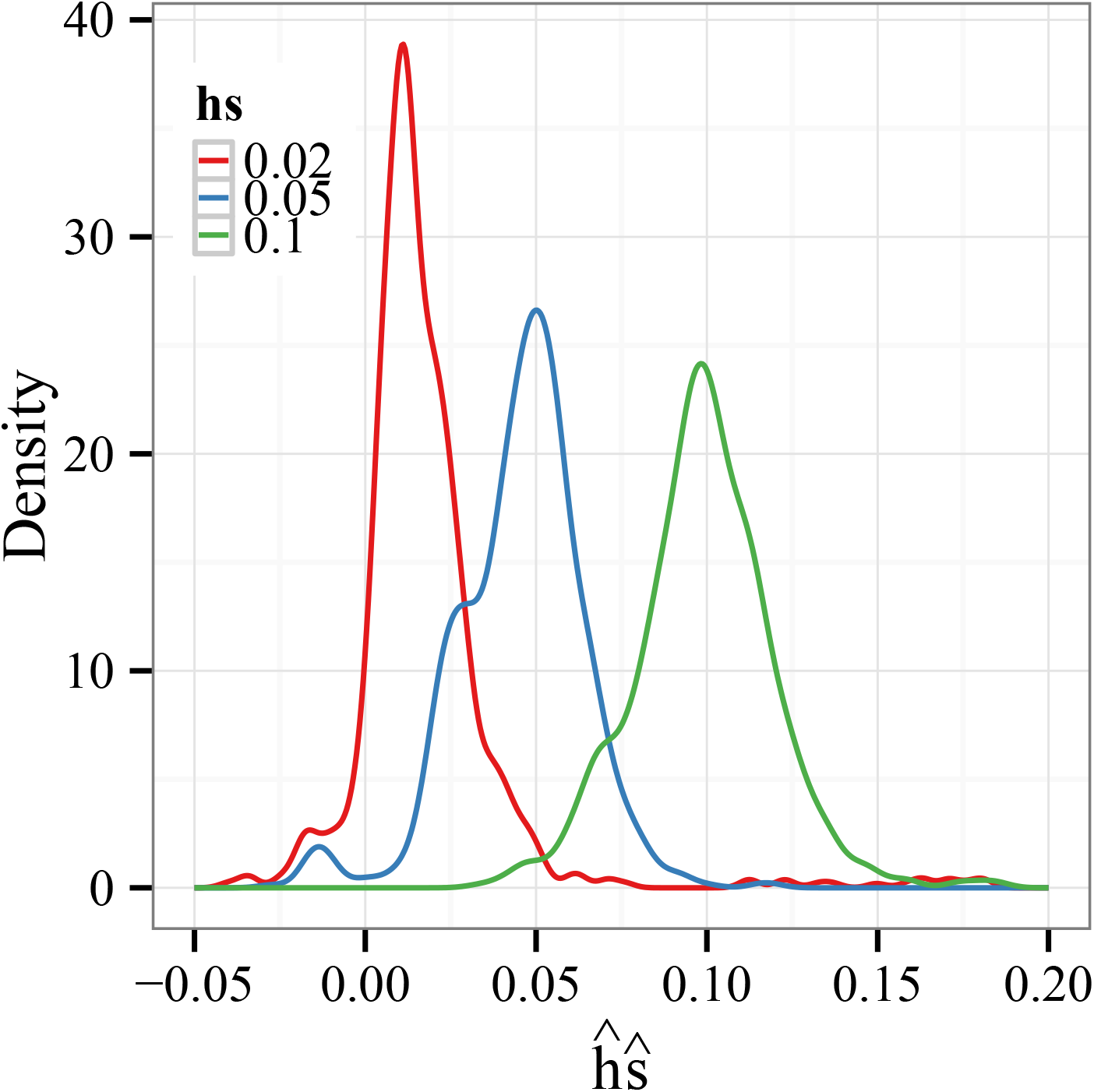
Overdominance estimation. Density estimates of the product *ĥ* · *ŝ* when the parameters are estimated jointly. The selection coefficient was fixed at *s* = 0.01 while the dominance parameter *h* was varied. In each simulation, the initial allelic frequency was restricted to lie outside the interval [0.4, 0.6] (see discussion in text). The mean of *ĥ* ·*ŝ* is quite close to the true value *hs* and the distribution is tightly concentrated around that value.

### Recombination rate estimation

Our multi-locus model can also be used to study phenomena which alter covariance between linked alleles. For example, in a region containing a recombination hotspot, covariance decreases markedly as increased recombination breaks down linkage disequilibrium. Using the same likelihood-based approach as above, the recombination rate within the hotspot can be estimated from E&R data. To test this, we simulated a region of length *L* = 100 kb in which the middle 2 kb region had an elevated recombination rate *r_H_* = *α · r*, where *r* = 10^−8^ is the background recombination rate and α ∈ {10, 10^2^, 10^3^}. For simplicity, we focused on the case of *C* = *∞* and assumed that the hotspot boundaries are known. For each simulation, a 30-locus model was fit using 10 randomly-selected loci from within the hotspot and 20 outside of it. Density estimates for the residual log_10_(*r̂H*).−.log_10_(*rH*) are shown in Figure 5. In all cases, the mode of the density occurs close to zero. A 3-order increase in the recombination rate is easily detected in experimental data, and a 2-order increase can also be estimated to well within an order of magnitude of accuracy. Increasing the recombination rate by only a factor of 10 leads to a fairly dispersed estimator, and it would be difficult to detect using the default experimental parameters.

**Figure 5.**
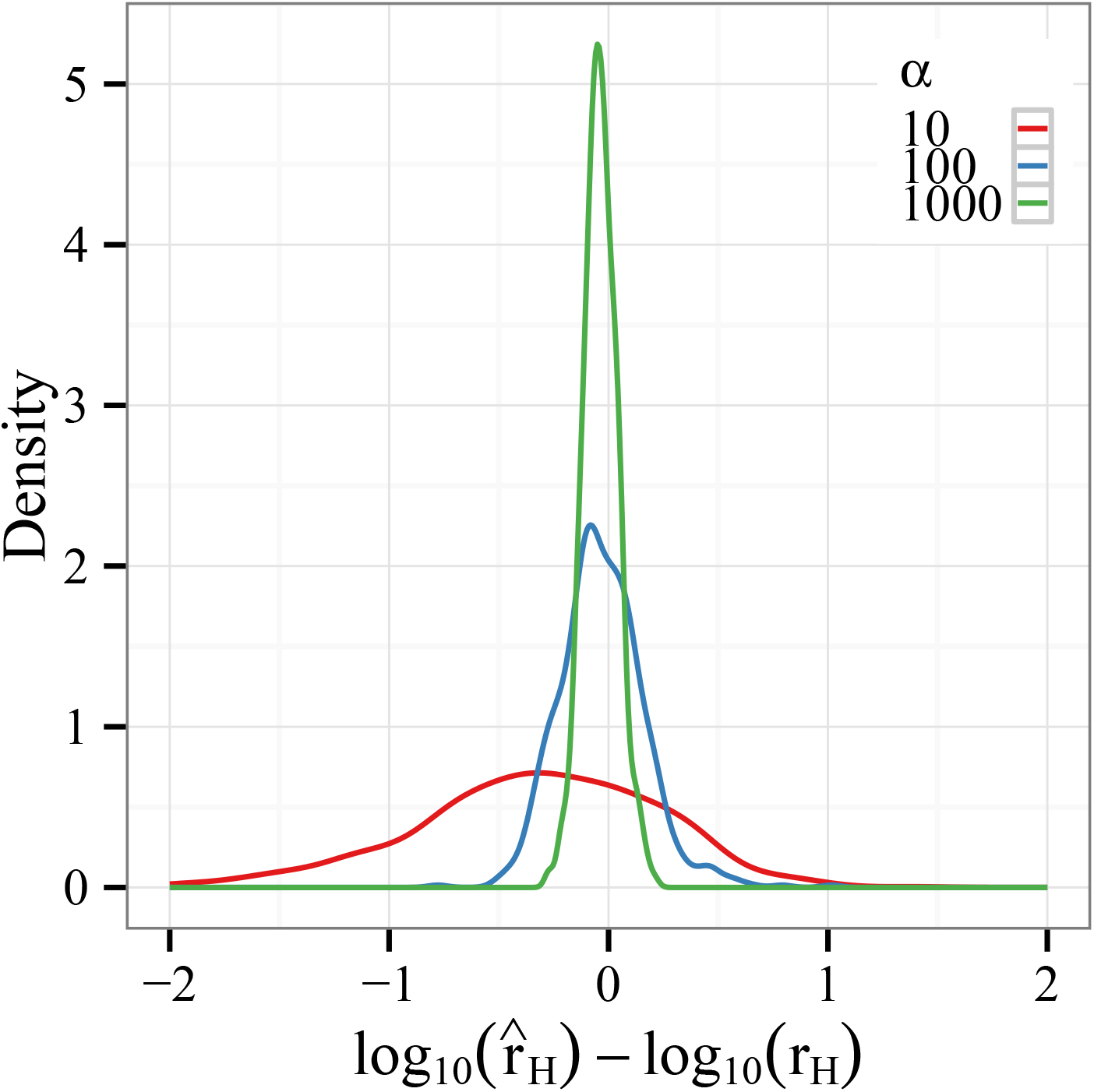
Hotspot estimation. A recombination hotspot was simulated by evolving a 100 kb region in which the recombination rate *r_H_* = *α · r* for the middle 2 kb (positions 49–51 kb) was increased by a multiplicative factor α ∈ {10, 100, 1000} above the baseline recombination rate *r*. The hotspot intensity *r̂H* was then estimated from experimental data. The figure shows density estimates of the residual log_10_(*r̂_H_*) − log_10_(*r_H_*) for each value of *α*.

### Effective population size estimation

As a final application of our method, we consider estimating the effective population size *N_e_* from experimental data. Up to now we have assumed that the (census) size *N* of the experimental population is fixed at a known value. In practice, the effective and census population sizes may differ due to various factors, including nonrandom mating and population structure. It could be interesting to quantify this effect by estimating *N_e_* in experimental data using the same likelihood-based procedures described above. Since our model approximates the Wright-Fisher process, in which *N_e_* = *N*, and simulations were carried out also assuming the Wright-Fisher model, we expect our estimate *N̂_e_* to be close to *N*. Figure 6 shows a scatter plot of *N̂_e_* versus *N* for 1,000 simulated E&R experiments. In each experiment, the population size *N* was chosen uniformly at random from the interval [10, 10^4^]. We see that the estimator is quite accurate for small population sizes and becomes more variable as *N* grows. This is expected since *N̂_e_* is essentially measuring genetic drift, which is of order *O*(1*/N)* as *N* grows. Thus, the inverse map taking drift to population size is well-conditioned for small *N* and becomes ill-conditioned as *N* grows.

**Figure 6.**
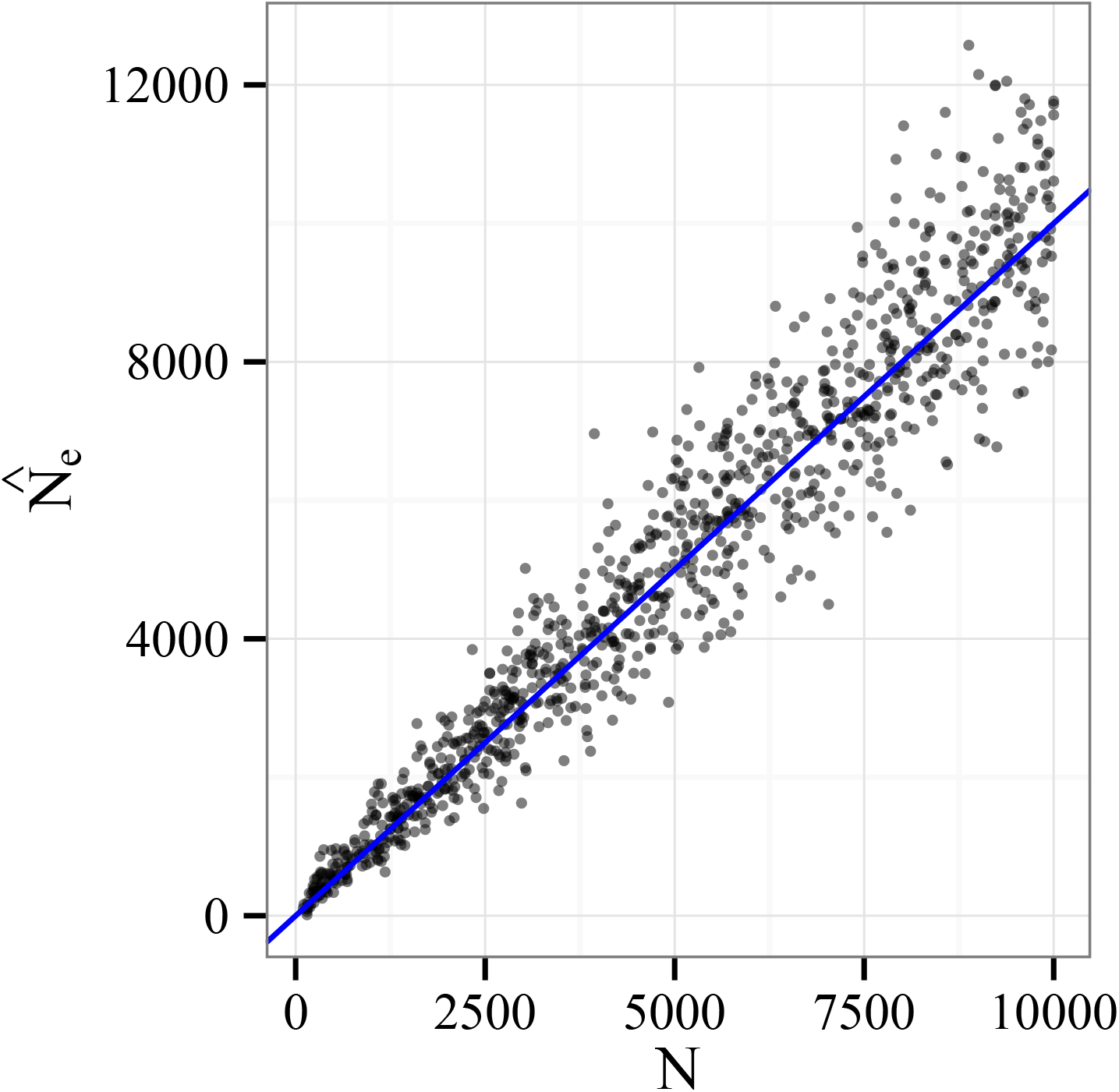
Effective population size estimation. The estimated effective population size (*N̂_e_*) versus the census population size (*N*) for 1,000 simulated E&R experiments. For each simulation, population size was chosen uniformly at random from the interval [10, 10^4^]. The estimator is quite accurate for small *N*, but becomes more variable as *N* grows. See text for discussion.

## Discussion

In this paper we have presented a model for analyzing time series data generated by evolve-and-resequence experiments. Our model is designed to analyze multiple recombining sites evolving in a moderately-sized population and potentially affected by measurement error. On data obtained from simulated E&R experiments combined with pooled sequencing, we have shown that it is possible to detect, localize and estimate the strength of selection in the range *s* ∈ [0.01, 0.10] in a population of moderate size (*N* ∼ 10^3^) and using a moderate number (*R* = 3) of experimental replicates. We have also explored the effect of the founding population composition (in terms of the number of founders) and sequencer effort (coverage depth, number of sampling time points, and time intervals between sampling) on the quality of these estimates. Finally, we have shown that our method can also be applied to study other phenomena of interest, including overdominance and effective population size; in particular, our work suggests that E&R data can be used to estimate recombination rates in putative hotspots in model organisms inferred by previous studies [5, 35, 36]. Space and time considerations have necessarily prevented us from considering many other combinations of experimental parameters which could be informative when designing E&R experiments. To enable other researchers to explore these options, we will make the computer code used in this study publicly available.

Experience has shown that the running time of our model is dominated by the recursive procedure used to calculate covariances between pairs of sites (see **Methods**). Thus, to fit a *K*-locus model sampled at *T* time points has computational complexity of order *O*(*K*^2^*T* ^2^). When performing the large number of simulations needed to benchmark our model, this quadratic scaling in the model size *K* prevented us from fitting models jointly using many more sites. Since our results suggest that estimation precision can be improved (in particular, at low coverage) by exploiting linkage information between sites, it could make sense in practice to expend additional computation time in order to add more sites into the model.

It is interesting to compare our findings with existing results. Feder *et al.* [29] suggest that power to detect selection is maximized when (positively) selected alleles are sampled as they rise in frequency, but before they have fixed. By a simple modification of their argument, the expected strength of selection required for a mutation in our simulated E&R experiments to achieve frequency *x_f_* in *T* time periods is given by

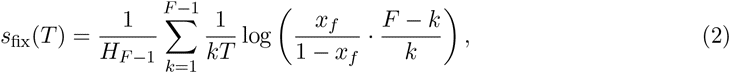

where 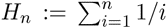 is the harmonic series. Above we generally chose *T* = 50 and *F* = 10; for *x_f_* = 0.95 we find that *s*_fix_(*T)* = 0.08 which roughly agrees with our finding (Figure 1) that medium and strong selection (*s* = 0.05, 0.1) could be reliably detected, while weaker selection was fairly difficult to detect. Our findings are somewhat more optimistic than those of Baldwin-Brown *et al.* [30], whose simulation results suggest that E&R experiments require a fairly large number of experimental replicates (*R* ≥ 25), founder haplotypes (*F ≥*500) and strong selection (*s ≥*0.1) in order to reliably detect and localize selected sites in a 1 Mb region. Since we used a smaller region for simulation (*L* = 100 kb), the results we report are not directly comparable; nevertheless, it is interesting that with many fewer replicates and haplotypes (*R* = 3 and *F* = 20) we could reliably detect the selected site in at least 50% of trials (Table 1). With sampled data the problem becomes harder, but we found that average coverage depth 30 still sufficed to discover the selected site from among the top four segregating sites in 50% of trials (Table 3).

Several extensions to our model could potentially be of use. For multi-locus estimation problems, our model requires that the haplotypic structure of the founding experimental population be known. In cases where this information is not known exactly, a Bayesian approach could be adopted in which model results are weighted by a prior on the space of initial haplotypic configurations. Such a procedure could allow the researcher to trade sequencing effort for computation time by decreasing the burden of initial sequencing that must be performed in order to establish the haplotypes of the founding lineages.

The other extreme of sequencing effort is to obtain haplotype data for a sample of individuals at each sampling generation, rather than to use pooled sequencing to infer only marginal allele frequencies. (Indeed, there is a discussion on the utility and power of pooled sequencing [37–40].) The same multi-locus model underlying our approach can be applied to develop a method for analyzing haplotypic time series data, and we will explore incorporating such an extension into our method.

Our approximation to the multi-locus Wright-Fisher process relies on a system of recursions which describe the evolution of neutral sites conditional on the presence of a linked selected site (see **Methods**). The process of generating those recursions has been automated [41] to handle more general scenarios including population structure and interaction between multiple selected sites. Our model could therefore be extended to handle these more complex scenarios at the expense of (potentially significantly) greater computational effort and data requirements.

## Methods

We consider the following model of an E&R experiment. A sexually reproducing population of *N* diploid individuals is evolved in discrete, non-overlapping generations. Pooled DNA sequencing [37] is performed *T* times at generations *t*_1_ < *t*_2_ < · · · < *t_T_*. If all sites are biallelic, the resulting dataset **D** ∈ [0, 1]^*T*×*L*×*R*^ counts relative frequency with which the *A*_1_ allele was observed for each combination of generation, locus and replicate. (The model is agnostic to which allele is called *A*_0_ or *A*_1_; interchanging the labels simply reverses the sign of the selection coefficient.)

Given **D** and a vector of underlying population-genetic parameters Θ, let 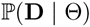 denote the model likelihood. In an idealized E&R experiment, generations are discrete and non-overlapping, mating is random, and the population size is fixed, so that likelihood is well approximated by the classical Wright-Fisher model of genome evolution:

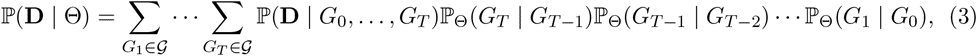

where 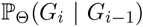 is the transition function of the discrete, many-locus Wright-Fisher Markov chain from genomic configuration *G_i−1_* to *G_i_* given parameters Θ, 𝒢 is the set of all possible genotypic configurations in a diploid population of size *N*, and 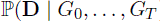) is the probability of the sequencer emitting **D** conditional on *G*_0_*,…, G_T_*. In our present formulation, we assume that *G*_0_ is known.

For typical problems, evaluating (3) is intractable since *|𝒢|* is very large and the transition density 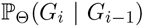 is difficult to compute and store. Asymptotic (i.e., diffusion) approximations to the transition density may be inaccurate if the population size *N* and/or scaled generation time 2*Nt* are small, as is common in an E&R experiment. Hence, alternative approximations to 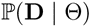 are needed to perform inference.

The approximation we make is as follows. Let X ≡ (*X_ijk_*) ∈ [0, 1]*^T^*^×*L*×*R*^ denote the (unobserved) population frequency of the *A*_1_ allele at each data point. Conditional on knowing **X**, and assuming that the DNA sequencer samples each site independently and binomially from the population, we have *D_ijk_ ∼* Binomial (*c_ijk_, X_ijk_*) where *c_ijk_* is the depth of sequencing coverage observed at this site. (Although sequencer coverage is random, we assume that it is independent of all other variables in the experiment and treat it as constant.) Marginalizing over the unobserved **X**, we obtain

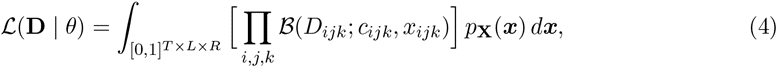

where 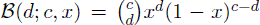 is the probability mass function of the binomial distribution and *p*X(*x*) is the density of **X**. Note that if each *c_ijk_* is large, as when the samples have been deeply sequenced, then the likelihood is (approximately) proportional to the density of **X**, i.e., ℒ(D|*θ*) ∝ *p_**X**_*(*x*), and the integral in (4) does not need to be evaluated. This computational savings can be useful when performing simulations.

To approximate the density *p***_X_**, we assume that, conditional on the initial genome configuration *G*_0_, the underlying allele frequencies *X_ijk_* are distributed according to a Gaussian process:

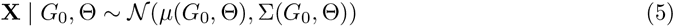

where, as described below, the first- and second-order moment functions *µ*(*·*) and Σ(*·*) are obtained by considering appropriate Wright-Fisher models. As described below, we are essentially approximating the complex joint distribution of allele frequencies by a sequence of simpler two-locus distributions. This approximation enables us to induce the correct mean and covariance structure in the random variable **X** while forgoing information captured in higher moments. Using this approximation we can perform tractable, likelihood-based inference while capturing salient aspects of the linkage-induced correlation present in the data.

### Moments of the Wright-Fisher process

To specify the model (5) we must compute the first-and second-order moments of **X** for any time *t_i_* ∈ {*t*_1_,…, *t_T_*}, locus *j* ∈ {1,…, *L*}, and replicate *k* ∈ {1,…, *R*}. Since the replicates are assumed to be independent and identically distributed, we suppress the dependence on index *k* for the remainder of this section.

The *L*-locus Wright-Fisher model with two alleles at each locus is a discrete-time Markov process 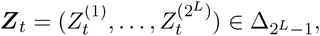 for *t* = 1, 2,…, where

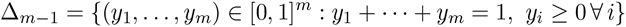

denotes an (*m* − 1)-dimensional simplex. The 2*^L^* different entries of ***Z****_t_* correspond to distinct haplotypes. For example, in a two-locus model with alleles *a, A* at the first locus and alleles *b, B* at the second locus, ***Z****_t_* is a 4-tuple with the entries corresponding to the population-wide fraction of *A*_1_*A*_1_*, A*_1_*A*_0_*, A*_0_*A*_1_, and *A*_0_*A*_0_ haplotypes.

Corresponding to the process ***Z****_t_* is the *L*-dimensional *marginal process* 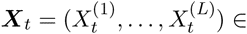 [0, 1]^*L*^ in which *X*^(^*^j^*^)^ denotes the population frequency of the *A*_1_ allele at locus *j* and time *t.* Thus, in the above two-locus example, if ***Z****_t_* = (0.1, 0.2, 0.3, 0.4) then ***X****_t_* = (0.3, 0.4) gives the population-wide marginal frequencies of the *A*_1_ alleles. It is this marginal process which we observe in a pooled sequencing experiment.

Since each entry of ***X****_t_* is a linear combination of the entries of ***Z****_t_*, it suffices to compute moments of the form 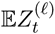 and 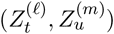 for arbitrary times *t, u* and loci *ℓ, m* There are two cases. In the neutral case (*s* = 0) we derive an analytic approximation to the covariance trajectory of ***Z****_t_*. In the case of fitness differences between genotypes (*s* ≠ 0), a different approximation is necessary. To calculate the moments in this case we perform a Taylor expansion of the process transition function about the drift-free (deterministic) sample path. This yields a system of recursions which can be used to solve numerically for the relevant moments in arbitrary generations. This approach was previously employed by Barton *et al.* [42] to obtain order *O*(1*/N)* approximations to these moments. Here we have used the same idea but automated the symbolic algebra and code generation needed to generate the recursions to higher orders of accuracy.

### Neutral case

In the case of neutrality, it suffices to consider covariances between pairs of sites in a two-locus haploid model. The one-generation transition function of the neutral two-locus Wright-Fisher model with recombination fraction *r* is

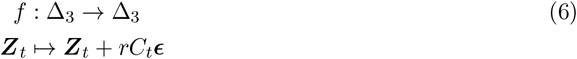

where *ϵ ≡* (−1, 1, 1, −1) and 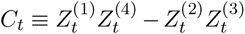 is the linkage disequilibrium at time *t*. Thus, conditional on ***Z****_t_* we have that 2*N ×* ***Z****_t_*_+1_ is multinomially distributed according to *f* (***Z****_t_*):

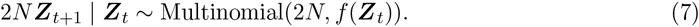

Using equation (7), we can derive an accurate approximation to the evolution of the covariance of the ***Z****_t_* process. In what follows we let *π* = (*z*^(1)^*, z*^(2)^*, z*^(3)^*, z*^(4)^) and *c*_0_ = *z*^(1)^*z*^(4)^ −*z*^(2)^*z*^(3)^ denote the initial distribution and linkage disequilibrium of the Wright-Fisher process under consideration.

#### Lemma 1.

To order 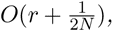

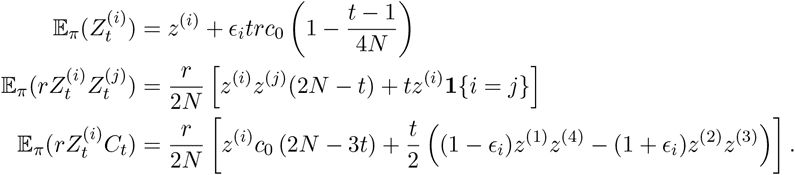

*Proof of Lemma 1*. By direct computation using the moment generating function of the multinomial distribution, we find that

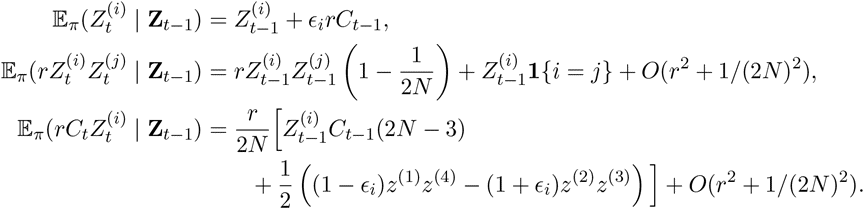

The results now follow by induction.

#### Corollary 2.

*To order* 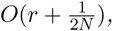

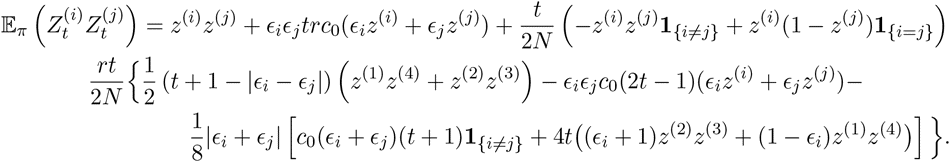

*Proof of Corollary 2.* This result is obtained by considering the conditional expectation 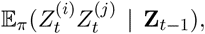 inducting on *t*, and checking cases for *i* and *j*. We illustrate the proof for the case *i* = *j* = 1 and omit the lengthy but routine computations used to check the remaining cases. (A Mathematica notebook which checks all cases is available from the authors upon request.) With *i* = *j* = 1, we have

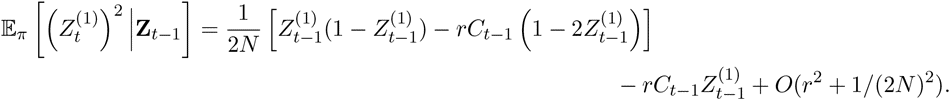

This yields the claim for *t* = 1. Taking expectation and applying the preceding lemma, we find that

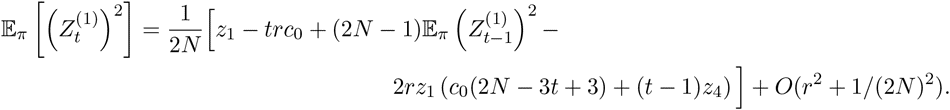

Applying the inductive hypothesis, we obtain

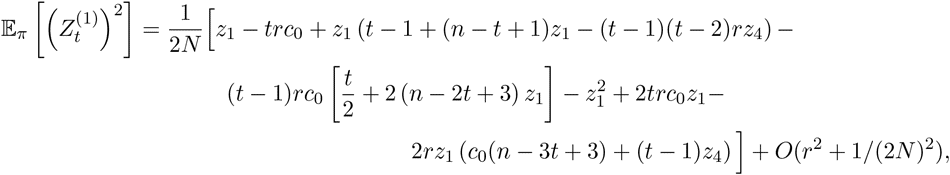

which agrees with the claim after some simplification.

The above results can be combined to give an 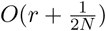 approximation to the within-generation covariance 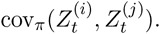. Using the same approach, we can also approximate the covariance between generations. Indeed, by Lemma 1 and the Markov property,

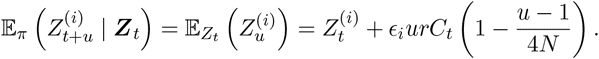

Hence,

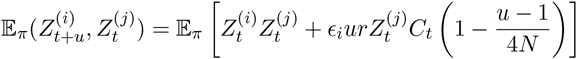

and each of the expectations on the right-hand side is given to order 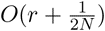 by the preceding results.

*Remark.* The constants subsumed in the 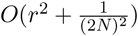 terms in the above expressions increase as *t* increases; in particular, we would not expect the approximation to be accurate if *tr* ∈ *O*(1). For our application typically *t* ≪ 1/*r*

### Non-neutral case

In the non-neutral case the transition operator *f* (***Z****_t_*) is a rational function of its arguments, so moments of ***Z****_t_*_+1_ depend on all moments all orders of ***Z****_t_*. A different form of approximation is needed in this case. Formally, we decompose ***Z****_t_* as 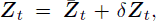 where 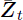 equals the deterministic trajectory in the absence of genetic drift, and *δ**Z**_t_* is a random disturbance away from the deterministic path. This permits a Taylor expansion of the relevant moments about the deterministic sample path, which yields a recursion for computing these moments in terms of lower order moments and moments from previous time periods.

To illustrate the method on a simple example, consider a one-locus Wright-Fisher model with diploid selection and no mutation. The relative fitnesses of *A*_0_*/A*_0_ and *A*_1_*/A*_1_ homozygote genotypes are given by 1 and 1 + *s*, respectively, whereas the relative fitness of the *A*_0_*/A*_1_ heterozygote is 1 + *hs*. The frequency of the *A*_1_ allele at time *t* is denoted *X_t_*. Conditional on *X_t_*, 2*N × X_t_*_+1_ has a binomial distribution with 2*N* trials and success parameter *f* (*X_t_*) [43], where

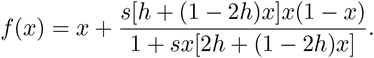

𝔼(*X_t_*_+1_) can be approximated by the Taylor expansion of *f* about the deterministic path 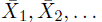:

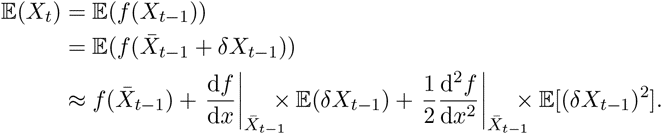

(For brevity we illustrate the method using expansions truncated to second order, however we have written software which automates these calculations to arbitrary order.) We also have 𝔼 (*X_t_*) = 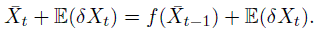. Combining these two equations yields

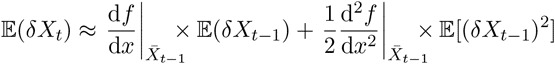

which is a recursion for computing 𝔼(*δX*_*t*_) in terms of the moments of *δX_t-_*_1_. Inductively assuming that we can compute 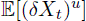 for *u* = 1; 2, we can compute 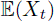 and var(*X_t_*) = var(*δX_t_*).

### Multi-locus case

The above idea can be extended to multiple loci in a straightforward manner. Recall ***Z****_t_* = 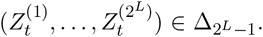. Conditional on ***Z_t_***, the vector 2*N × Z_t+1_* is multinomially distributed with success probabilities *f* (***Z****_t_*). The form of 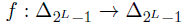 varies according to the underlying model; we describe our choice of *f* in the following section.

As in the one-locus case, write 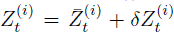 where 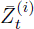 is the deterministic trajectory which would be followed by 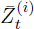 if the population were infinite, and *δ*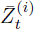 is a random disturbance.

(Note that in general, 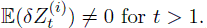 For *u, v* non-negative integers, we have

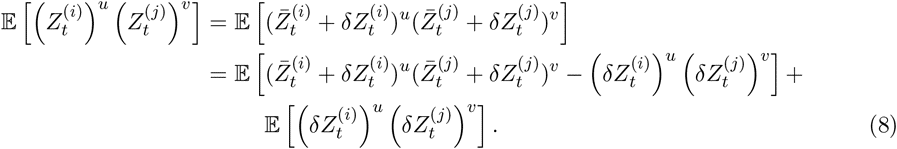

From the conditional distribution 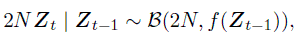 we have

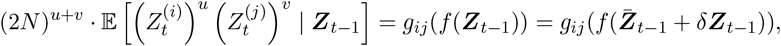

where 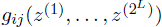 is a polynomial in 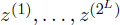 which can be computed using the moment generating function of the multinomial distribution. By performing a Taylor expansion of *ϕ_ij_* ≡ *g_ij_ ∘ f* about the deterministic path 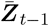 and taking expectations, we get another formula for 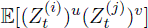 in terms of moments of 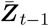

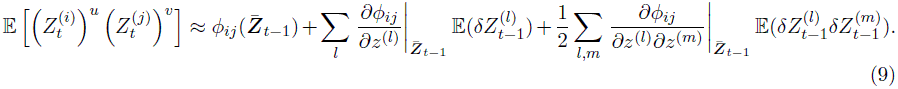

For *u* + *v ≤* 2, comparing (8) and (9) yields a recursion for computing 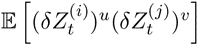 terms of moments of *δ**Z**_t_* of total degree strictly less than *u* + *v*, and moments *δ**Z**_t−1_* of total degree at most *u* + *v*. The latter feature is important for computation because it implies that we only need to compute a bounded number of terms in each recursive step, which would not be the case if we had instead expanded the function *ϕ_ij_*(*·*) about zero with respect to model parameters (for example, selection or mutation).

The recursive nature of the above algorithm lends itself to computing moments of the form 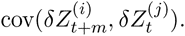. Stopping the recursion m time steps into the past, we obtain an expression of the form 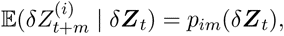 where 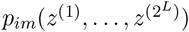 is a polynomial. Hence,

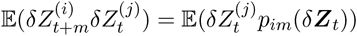

is again a recursion involving moments of *δ**Z**_t_* which can be solved using the techniques described above.

### Transition Function

The transition function *f* described in the previous section describes the effects of recombination, selection and, optionally, mutation on gamete formation from one generation to the next. Since mutation is rare for the population sizes and time scales typical of E&R experiments on macroscopic organisms, we do not treat it in our model. The one-and two-locus moments studied in the previous section will be altered by the presence of a linked site which is under selection. Since we assume that within the region of interest at most one segregating site has a nonzero selection coefficient, it is appropriate to employ a three-locus model which describes the forward evolution of one selected and two neutral loci. The exact form of this model depends on location of the selected locus (i.e., whether it is between the neutral loci or to one side of them) and has been derived by Stephan *et al.* [44] using the general framework of Kirkpatrick *et al.* [41]. We implemented these functions in the symbolic algebra program *Maple* in order to automate the recursive Taylor expansion procedure described above.

### Simulation

Our procedure for simulating an E&R experiment was the following. To generate realistic patterns of standing variation, a set of *F* founder lines was sampled from the coalescent with recombination using the program `ms` [45]. Recombination and mutation rates and the effective population size were set to biologically plausible values for *D. melanogaster*, a common model organism used in E&R studies (*r* = 2 *×* 10^−8^/bp/gen, *µ* = 10*^-^*^9^/bp/gen, *N* = 10^6^) [46]. Each founder line was cloned 2*N/F* times to generate an initial diploid population of size *N*. This replication step is intended to mimic the practice using of (nearly-)homozygous recombinant inbred founder lines to initialize an E&R experiment. Next, the experimental population of size *N* was simulated forward in time using the discrete-time simulator `simuPOP` [47]. Finally, alleles were sampled binomially and independently at each locus and time point to simulate next-generation sequencing. Parameters for the forward simulation and sampling were varied from scenario to scenario as described in the main text. The output of the simulation consisted of the haplotypes of the initial founder lines and the frequency of each segregating site (potentially after sampling) at each time point. All simulations were performed on a machine with 2 *×* 2.5 GHz AMD Opteron 6380 processors (32 cores total) and 256 GB of memory.

## Acknowledgments

The authors thank Julien Ayroles, Anand Bhaskar, Andy Clark, Graham Coop, Tony Long, Christian Schlötterer, and Matthias Steinrücken for helpful comments and discussions. This research is supported in part by an NIH National Research Service Award Trainee appointment on T32-HG00047, an NIH Grant R01-GM094402, and a Packard Fellowship for Science and Engineering.

